# Genetic Influences on Neural Responses in Placebo Analgesia Circuitry

**DOI:** 10.64898/2025.12.09.693210

**Authors:** Jörgen Rosén, Gránit Kastrati, Fredrik Åhs, Karin Jensen

**Affiliations:** Department of Clinical Neuroscience, Karolinska Institutet, Stockholm, Sweden; Department of Psychology, Mid Sweden University, Östersund, Sweden

**Keywords:** fMRI, heritability, placebo analgesia, pain, translational

## Abstract

Placebo analgesia is a well-established medical phenomenon with overlapping neural representations between humans and rodents, but the genetic contribution to this conserved neural circuitry remains unknown. Using functional brain imaging in 305 monozygotic and dizygotic twins, we quantified the heritability of neural activation in placebo analgesia brain circuity during evoked pain, including the rostral anterior cingulate cortex, brainstem and cerebellum, using the ACE model. Neural responses showed significant heritability, with additive genetics accounting for 21 to 49% of the variance (A = 0.21-0.49), indicating that genetics have a moderate effect on pain-evoked brain activity within the placebo analgesia circuitry. For the first time, our findings reveal a heritable component of the neural network supporting placebo analgesia. By linking genetic variations to neural responses in placebo analgesia circuitry, this study bridges human and preclinical research and opens new ways for improving clinical trial methodology and tailoring pain therapies to individual genetic profiles.

## Introduction

Placebo analgesia is a well-documented phenomenon in humans ^1^, in which expectations of pain relief can lead to measurable hypoalgesia ^2^. Even if the placebo effect is commonly observed in the placebo arms of randomized controlled trials, placebo effects extend beyond the context of clinical trials. They are embedded within the therapeutic effects of all active treatments as a distinct placebo-component that shape the overall clinical outcomes ^3^ and thus have broad implications for all healthcare ^4^.

The first experiment to elucidate the biological mechanisms of placebo analgesia ^5^ used an acute pain model of third molar extraction and found that placebo-induced pain relief was mediated by release of endogenous opioids. This conclusion was supported by the finding that placebo analgesia was reversed by administration of an opioid antagonist (naloxone). By showing that the placebo effect engaged a specific neurobiological pathway, the study demonstrated that placebo analgesia is not merely psychological but rooted in identifiable physiological mechanisms. The advent of non-invasive brain imaging techniques has, since then, provided detailed anatomical evidence for the recruitment of an opioid-enriched inhibitory pathway during placebo analgesia, including the rostral Anterior Cingulate Cortex (rACC) and the brainstem ^6–8^, including specific somatotopic responses in the periaqueductal gray (PAG) ^9^ and evidence for inhibition at the level of the spinal cord ^10^. The use of naloxone during functional Magnetic Resonance Imaging (fMRI) ^7^ led to a disruption of the rACC-brainstem connectivity, and reduced placebo responses, suggesting that endogenous activation of this specific opioid pathway is central for placebo analgesia.

Although placebo effects have been well-established in humans, translation to animal models has been limited. Two pioneering studies used behavioral models of placebo analgesia in mice^11,12^ and provided more detail to the previous evidence for endogenous activation of opioid-enhanced inhibitory circuitry in humans. Chen et al. ^12^ developed a behavioral assay for placebo analgesia in mice and identified a neural pathway mediating the analgesic effect: rACC neurons projecting to the pontine nucleus in the brainstem. Using calcium imaging, electrophysiology, and transcriptomics, they showed that manipulation of this pathway has a causal role for achieving placebo analgesia. As a new addition to the previously described placebo analgesia brain circuitry, cerebellar Purkinje cells were found to mirror the rACC-brainstem activity in mice ^12^, implicating a previously unknown brain circuit for endogenous pain modulation in *humans*.

The cerebellum has previously been described in a neuroimaging meta-analysis of placebo analgesia ^13^ and a lesion study suggests that placebo analgesia is reduced in patients with cerebellar infarction due to impaired pain inhibitory control ^14^. Yet, the cerebellum has not been recognized as an influential component of the pain inhibitory brain circuit in humans, and its potential role in human placebo analgesia is unclear.

While previous research has identified specific neural substrates involved in placebo analgesia, the role of genetic factors in shaping these neural responses has remained largely unexplored. In this study, we present the first investigation to examine the heritability of the placebo analgesia circuitry using a twin fMRI design. By recruiting a large sample of monozygotic and dizygotic twins (n = 305) and applying functional neuroimaging, we aimed to disentangle genetic and environmental influences on pain-evoked neural responses in placebo brain regions recently identified in humans and rodents^12^, i.e., the rostral anterior cingulate cortex (rACC), brainstem and cerebellum. This novel approach provides valuable insights into the genetic basis of placebo responsiveness and endogenous pain modulation, offering a new framework for understanding individual differences in expectancy-driven pain relief.

## Results

### Brain activations

Significant pain-evoked activations were found in multiple brain regions associated with pain processing and modulation. Table 1 highlights robust activation within several subregions of the anterior cingulate cortex (ACC), including right and left Area 33, p32, p24c, and p24ab, as well as the rostral ACC (rACC). Additionally, cerebellar nuclei such as the interposed, dentate, and fastigial nuclei showed consistent involvement. Other regions with statistically significant activation (p < 0.05, Bonferroni corrected) during evoked pain include multiple areas of the insula (Id3, Id5, Ig2, Id1, Ia3, Id6, Ia1, Id9, Ig1, Id2, Id4, Id8, Id10, Ig3, Ia2), the inferior parietal lobule (PFcm, PFop, PFt, PFm, PF), frontal operculum (Op5, Op6, Op7, Op8, Op9), postcentral gyrus (3a, 3b, 1, 2), occipital cortex (hOc1, hOc2, hOc3d, hOc4d, hOc6), amygdala (LB, CM), metathalamus (CGL, CGM), superior parietal lobule (5L, 5Ci, 7M, 7PC, 7P), temporal regions (TE 1 0, TE 1 1, TE 2 1, TE 2 2, TE 3), fusiform gyrus (FG1, FG3), orbitofrontal cortex (Fo6, Fo7, Fo3), and basal forebrain (Ch 4). The statistically significant t-values and confidence intervals for these regions further support the widespread and coordinated engagement of these areas during painful stimuli, emphasizing their contribution to both the sensory and modulatory aspects of pain.

**Table 1.** Brain regions activated during painful stimulation. The table presents all brain regions that exhibited statistically significant activation (p < 0.05, Bonferroni corrected) in response to painful stimuli, as identified using the Julich brain atlas and parcellated brainstem regions derived from a probability atlas based on the Human Connectome Project.

| **Brain region** | ***t*-value (*n* =**<br>**264)** | **95% *CI***<br>***low*** | **95% *CI***<br>***high*** |
| --- | --- | --- | --- |
| Area Id3 (Insula) right | 18.15 | 0.43 | 0.53 |
| Area Id5 (Insula) right | 18.09 | 0.40 | 0.50 |
| Area Ig2 (Insula) right | 16.38 | 0.35 | 0.45 |
| Area PFcm (IPL) right | 16.17 | 0.43 | 0.55 |
| Area Id3 (Insula) left | 16.05 | 0.34 | 0.44 |
| Area Id1 (Insula) right | 15.89 | 0.31 | 0.40 |
| Area Ia3 (Insula) right | 15.77 | 0.90 | 1.16 |
| Area Id6 (Insula) right | 15.71 | 0.30 | 0.39 |
| Area Ig2 (Insula) left | 15.66 | 0.31 | 0.40 |
| Area PFop (IPL) right | 15.63 | 0.44 | 0.57 |
| Area Ia1 (Insula) right | 15.60 | 0.38 | 0.49 |
| Area Ia1 (Insula) left | 15.43 | 0.38 | 0.49 |
| Area Id1 (Insula) left | 15.05 | 0.40 | 0.53 |
| Area OP3 (POperc) right | 14.99 | 0.48 | 0.62 |
| Area OP4 (POperc) right | 14.65 | 0.48 | 0.62 |
| Area Op6 (Frontal Operculum) right | 14.43 | 0.69 | 0.91 |
| Area Id9 (Insula) right | 14.34 | 0.34 | 0.45 |
| Area PFt (IPL) right | 14.05 | 0.23 | 0.31 |
| Area Ia3 (Insula) left | 14.04 | 0.35 | 0.46 |
| Area OP1 (POperc) right | 13.95 | 0.52 | 0.69 |
| Area Id5 (Insula) left | 13.92 | 0.34 | 0.45 |
| Area OP2 (POperc) right | 13.74 | 0.40 | 0.54 |
| Area Id2 (Insula) right | 13.58 | 0.37 | 0.50 |
| Area Op5 (Frontal Operculum) right | 13.49 | 0.37 | 0.50 |
| Area PFop (IPL) left | 13.29 | 0.41 | 0.55 |
| Area hOc1 (V1 17 CalcS) left | 12.11 | 0.43 | 0.60 |
| Area 44 (IFG) right | 11.88 | 0.25 | 0.35 |
| Area Id6 (Insula) left | 11.68 | 0.22 | 0.31 |
| Area OP4 (POperc) left | 11.39 | 0.29 | 0.41 |
| Area OP3 (POperc) left | 11.36 | 0.34 | 0.48 |
| Area PF (IPL) left | 11.32 | 0.22 | 0.31 |
| Area Ig1 (Insula) right | 11.13 | 0.23 | 0.33 |
| Frontal to Temporal I (GapMap) right | 11.12 | 0.27 | 0.38 |
| Area OP2 (POperc) left | 11.01 | 0.35 | 0.50 |
| Area Op8 (Frontal Operculum) right | 10.98 | 0.25 | 0.36 |
| Area Op7 (Frontal Operculum) right | 10.82 | 0.40 | 0.57 |
| Area Id2 (Insula) left | 10.41 | 0.20 | 0.29 |
| Area 45 (IFG) right | 10.39 | 0.21 | 0.30 |
| Area 5L (SPL) right | 10.36 | 0.24 | 0.35 |
| Area PF (IPL) right | 10.32 | 0.24 | 0.36 |
| Area hOc6 (POS) left | 10.28 | 0.34 | 0.49 |
| Area PFm (IPL) left | 10.20 | 0.22 | 0.33 |
| Area Op6 (Frontal Operculum) left | 10.17 | 0.26 | 0.39 |
| Area Ig1 (Insula) left | 10.12 | 0.24 | 0.35 |
| Area 33 (ACC) right | 10.03 | 0.16 | 0.24 |
| Area Id4 (Insula) right | 9.92 | 0.21 | 0.31 |
| Area hOc2 (V2 18) right | 9.91 | 0.31 | 0.46 |
| Area PFm (IPL) right | 9.89 | 0.19 | 0.29 |
| Area Id9 (Insula) left | 9.85 | 0.23 | 0.34 |
| Area hOc3d (Cuneus) left | 9.73 | 0.29 | 0.43 |
| Area PFcm (IPL) left | 9.71 | 0.28 | 0.42 |
| Area hOc1 (V1 17 CalcS) right | 9.58 | 0.27 | 0.42 |
| Area 5Ci (SPL) left | 9.54 | 0.19 | 0.29 |
| Frontal to Temporal II (GapMap) right | 9.30 | 0.12 | 0.19 |
| Area Op7 (Frontal Operculum) left | 9.28 | 0.20 | 0.31 |
| Frontal to Occipital (GapMap) left | 9.14 | 0.13 | 0.20 |
| Area hOc2 (V2 18) left | 9.11 | 0.29 | 0.45 |
| Area 5Ci (SPL) right | 8.74 | 0.16 | 0.25 |
| Area Ig3 (Insula) right | 8.73 | 0.21 | 0.34 |
| CGL (Metathalamus) right | 8.60 | 0.23 | 0.36 |
| Area Ph1 (PhG) left | 8.40 | 0.17 | 0.27 |
| Area 45 (IFG) left | 8.35 | 0.17 | 0.28 |
| Area 7M (SPL) left | 8.23 | 0.38 | 0.61 |
| Area Id8 (Insula) right | 8.22 | 0.17 | 0.28 |
| Area Fo7 (OFC) left | 8.09 | 0.16 | 0.27 |
| Area p32 (pACC) left | 8.06 | 0.15 | 0.24 |
| Area hPO1 (POS) left | 7.99 | 0.20 | 0.33 |
| Frontal to Occipital (GapMap) right | 7.90 | 0.11 | 0.19 |
| Area 5M (SPL) left | 7.88 | 0.17 | 0.29 |
| Area Ph3 (PhG) right | 7.70 | 0.14 | 0.23 |
| CGL (Metathalamus) left | 7.65 | 0.14 | 0.24 |
| Area hOc4d (Cuneus) left | 7.64 | 0.20 | 0.34 |
| Area Id10 (Insula) left | 7.60 | 0.12 | 0.21 |
| Area Id10 (Insula) right | 7.59 | 0.13 | 0.21 |
| Area Id4 (Insula) left | 7.52 | 0.13 | 0.23 |
| Area 5M (SPL) right | 7.51 | 0.14 | 0.24 |
| Area Op5 (Frontal Operculum) left | 7.51 | 0.17 | 0.30 |
| Area p24c (pACC) right | 7.45 | 0.11 | 0.19 |
| Area STS1 (STS) right | 7.42 | 0.16 | 0.28 |
| Area 7M (SPL) right | 7.34 | 0.28 | 0.49 |
| Area hOc3d (Cuneus) right | 7.25 | 0.17 | 0.30 |
| Area p24c (pACC) left | 7.20 | 0.13 | 0.23 |
| Area Ph3 (PhG) left | 7.19 | 0.14 | 0.25 |
| Area Op9 (Frontal Operculum) right | 7.17 | 0.17 | 0.30 |
| Area Op9 (Frontal Operculum) left | 7.16 | 0.17 | 0.30 |
| Area Ia2 (Insula) right | 7.12 | 0.29 | 0.51 |
| Frontal to Temporal I (GapMap) left | 6.98 | 0.14 | 0.25 |
| Area OP1 (POperc) left | 6.96 | 0.17 | 0.31 |
| Temporal to Parietal (GapMap) left | 6.94 | 0.07 | 0.13 |
| CGM (Metathalamus) right | 6.93 | 0.11 | 0.20 |
| Area TE 1 0 (HESCHL) left | 6.91 | 0.33 | 0.59 |
| Area 5L (SPL) left | 6.79 | 0.11 | 0.20 |
| Interposed Nucleus (Cerebellum) right | 6.79 | 0.09 | 0.17 |
| Area PGp (IPL) left | 6.68 | 0.10 | 0.19 |
| Area TE 2 1 (STG) right | 6.61 | 0.12 | 0.23 |
| Temporal to Parietal (GapMap) right | 6.60 | 0.08 | 0.14 |
| Area p32 (pACC) right | 6.58 | 0.11 | 0.20 |
| Area Ig3 (Insula) left | 6.49 | 0.11 | 0.20 |
| Area TPJ (STG SMG) left | 6.44 | 0.10 | 0.19 |
| Area FG1 (FusG) left | 6.34 | 0.14 | 0.27 |
| Frontal I (GapMa(p right | 6.15 | 0.07 | 0.13 |
| Area TE 3 (STG) right | 6.12 | 0.10 | 0.20 |
| Area Fo6 (OFC) right | 6.09 | 0.11 | 0.22 |
| Ventral Dentate Nucleus (Cerebellum) left | 6.03 | 0.06 | 0.12 |
| Area p24ab (pACC) right | 5.97 | 0.09 | 0.18 |
| LB (Amygdala) right | 5.95 | 0.09 | 0.17 |
| Area hOc4d (Cuneus) right | 5.85 | 0.13 | 0.26 |
| Area 44 (IFG) left | 5.82 | 0.08 | 0.17 |
| Ventral Dentate Nucleus (Cerebellum)<br>right | 5.76 | 0.06 | 0.12 |
| Area 3b (PostCG) right | 5.76 | 0.08 | 0.17 |
| Area 3a (PostCG) right | 5.75 | 0.12 | 0.24 |
| Area 1 (PostCG) right | 5.69 | 0.09 | 0.18 |
| Area hOc6 (POS) right | 5.67 | 0.09 | 0.19 |
| Area p24ab (pACC) left | 5.60 | 0.08 | 0.16 |
| Frontal I (GapMa(p left | 5.42 | 0.06 | 0.13 |
| Area 7PC (SPL) right | 5.42 | 0.08 | 0.17 |
| Area STS2 (STS) right | 5.27 | 0.06 | 0.14 |
| Area TE 1 0 (HESCHL) right | 5.26 | 0.10 | 0.23 |
| Area hPO1 (POS) right | 5.16 | 0.11 | 0.24 |
| CM (Amygdala) right | 5.02 | 0.18 | 0.41 |
| Area 7P (SPL) left | 5.02 | 0.09 | 0.22 |
| Area Id8 (Insula) left | 5.00 | 0.07 | 0.15 |
| Area TE 1 1 (HESCHL) left | 4.88 | 0.10 | 0.23 |
| CGM (Metathalamus) left | 4.84 | 0.06 | 0.14 |
| Area TE 1 1 (HESCHL) right | 4.82 | 0.05 | 0.13 |
| BST (Bed Nucleus) right | 4.82 | 0.05 | 0.11 |
| Area Fo7 (OFC) right | 4.75 | 0.07 | 0.16 |
| Ch 4 (Basal Forebrain) left | 4.63 | 0.07 | 0.17 |
| Ch 4 (Basal Forebrain) right | 4.55 | 0.07 | 0.18 |
| Area STS2 (STS) left | 4.51 | 0.05 | 0.12 |
| Area 8d1 (SFG) left | 4.51 | 0.04 | 0.10 |
| Area 8d1 (SFG) right | 4.51 | 0.04 | 0.09 |
| Area TE 2 2 (STG) left | 4.48 | 0.05 | 0.14 |
| Area TE 1 2 (HESCHL) right | 4.48 | 0.07 | 0.19 |
| Area Fo3 (OFC) right | 4.45 | 0.06 | 0.15 |
| Fastigial Nucleus (Cerebellum) right | 4.28 | 0.06 | 0.15 |
| Area 2 (PostCS) right | 4.26 | 0.04 | 0.11 |
| Area FG3 (FusG) left | 4.23 | 0.04 | 0.12 |
| Area Fp2 (FPole) right | 4.07 | 0.06 | 0.17 |
| Area Id7 (Insula) right | 4.00 | 0.05 | 0.16 |
| Area IFJ2 (IFS PreCS) right | 3.96 | 0.04 | 0.12 |
| Dorsal Dentate Nucleus (Cerebellum) left | 3.90 | 0.03 | 0.08 |
| Dorsal Dentate Nucleus (Cerebellum) right | 3.90 | 0.03 | 0.09 |
| Area STS1 (STS) left | 3.90 | 0.04 | 0.12 |
| HC Subiculum (Hippocampus) right | 3.83 | 0.05 | 0.15 |

**Figure 1.**
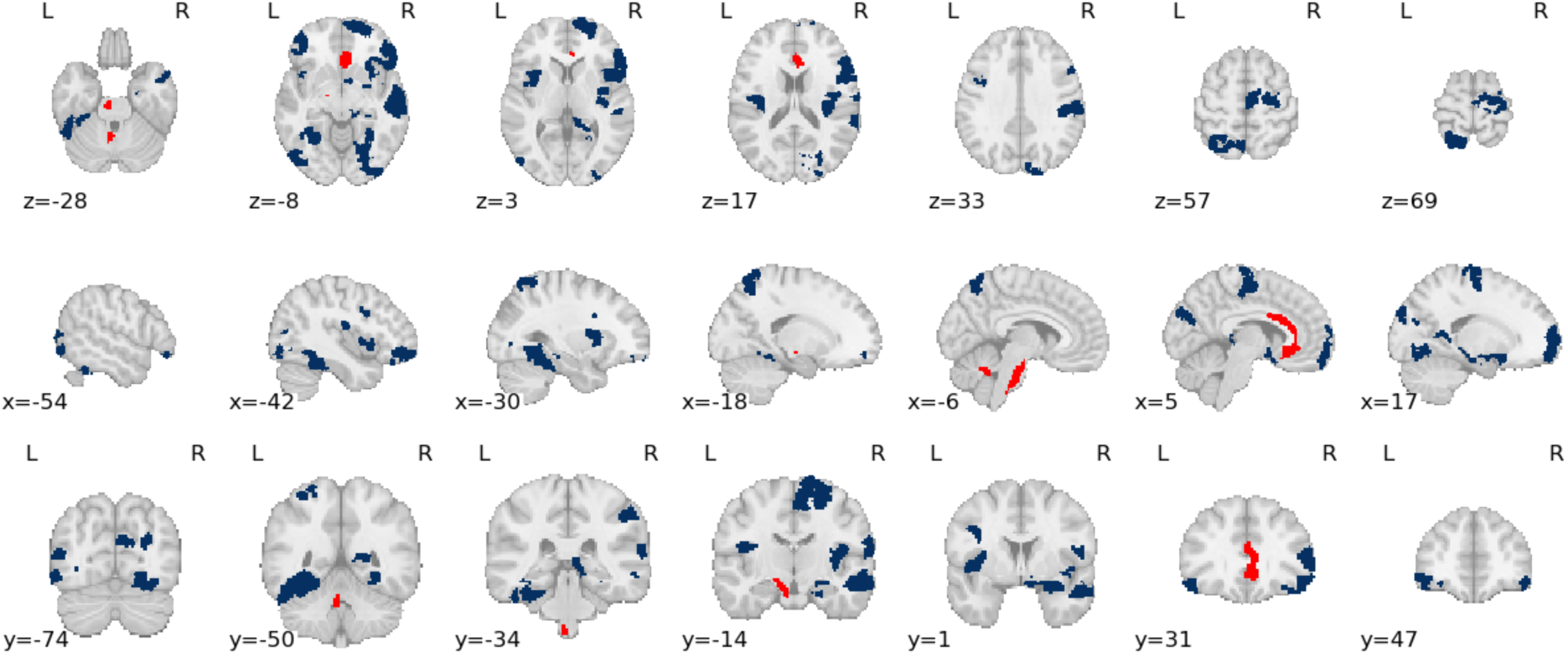
Heritability of pain-evoked brain activations in the placebo analgesia brain circuitry. Images display brain regions with significant additive heritability during evoked pain. Significant brain regions within the pre-defined placebo analgesia circuitry in *red*, all other brain regions with significant additive heritability in *blue*.

### Genetic influence on neural responses to pain

The analysis of heritability in neural responses to pain, as detailed in Table 2, demonstrates that several brain regions show significant additive genetic influence (A) during evoked pain. Notably, within the pre-defined placebo analgesia network, highlighted in red, in Figure 2, regions such as the left brainstem (pons), rostral anterior cingulate cortex (rACC), and cerebellar nuclei exhibited significant heritability estimates with a moderate rate of heritable influence. For example, the additive heritability (A) reached 0.45 in the left brainstem, 0.32 in the rACC, and up to 0.29 in the cerebellum, indicating that genetic factors account for a substantial portion of the variance in neural responses within these key nodes of the placebo analgesia circuitry. These findings suggest that the neural substrates underlying placebo analgesia are shaped in part by inherited genetic factors, particularly in the brainstem, rACC, and cerebellar regions. Other regions with significant heritability included the insula, inferior parietal lobule, frontal operculum, and fusiform gyrus.

**Figure 2.**
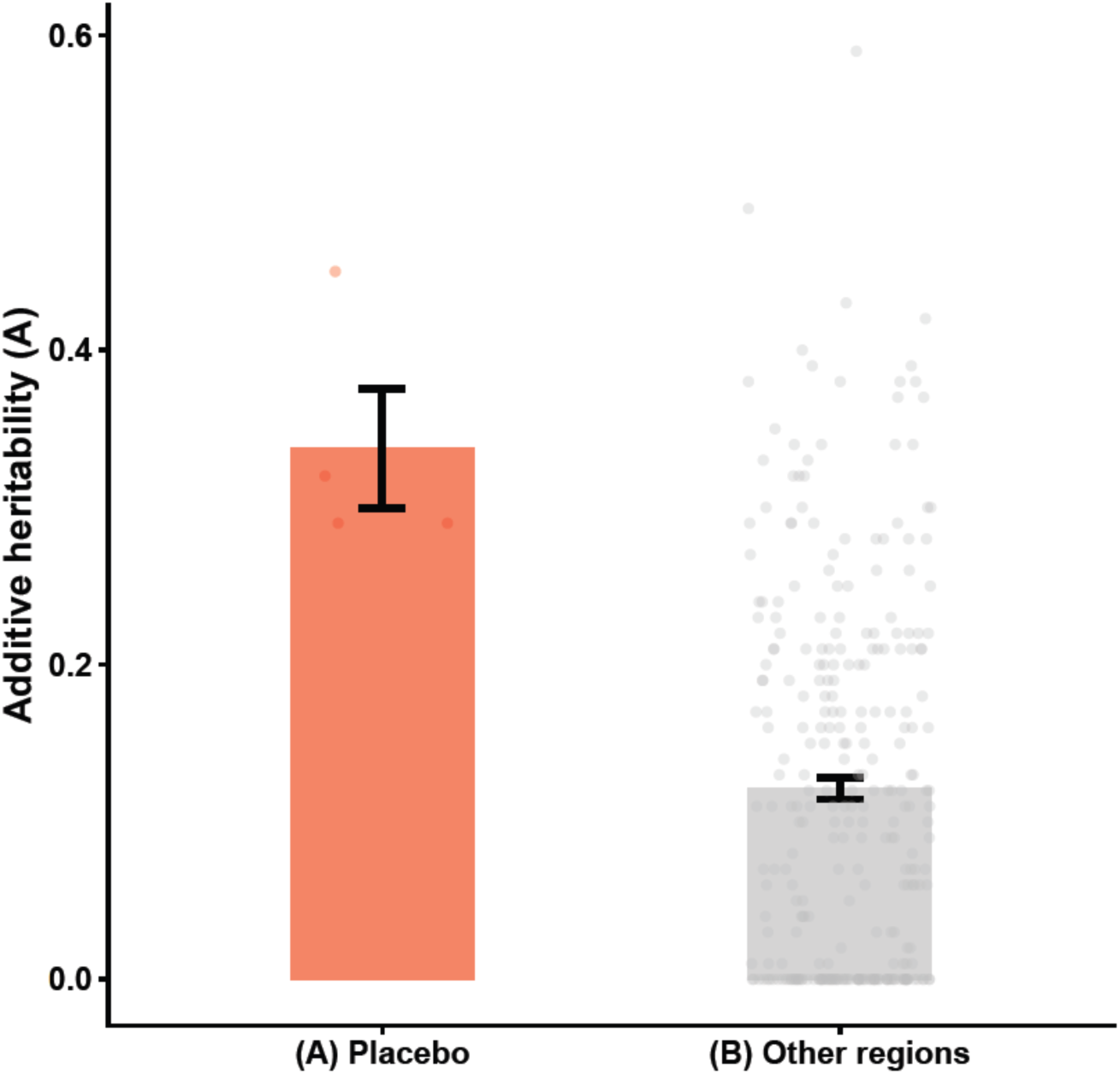
Sensitivity of brain regions with significant additive heritability. Bar charts represent mean values for (A) brain regions within the pre-defined placebo analgesia circuitry with statistically significant additive heritability (p < 0.05 Bonferroni corrected), and (B) estimates for the rest of the brain. Error bars denote standard deviation. Brain regions were identified using the Julich Brain Atlas, as well as brainstem regions delineated through an atlas based on the Human Connectome Project.

**Table 2.** Heritability of pain-evoked brain activations. Additive heritability (A), common environment (C), and unique environment effects on neural responses to evoked pain are shown for brain regions with statistically significant additive heritability (p < 0.05, Bonferroni corrected). Regions in **bold** belong to the pre-defined placebo analgesia network. Masks are annotated according to the Julich brain atlas and include parcellated brainstem regions derived from the Human Connectome Project.

| **Brain region** | ***MZ-***<br>***pairs*** | ***DZ-***<br>***pairs*** | ***MZ***<br>***corr*** | ***DZ***<br>***corr*** | ***A*** | ***C*** | ***E*** |
| --- | --- | --- | --- | --- | --- | --- | --- |
| Area IFJ2 (IFS PreCS) left | 59 | 66 | 0.49 | 0.24 | 0.49 | 0.00 | 0.51 |
| **Brainstem pons L** | 46 | 57 | 0.45 | 0.22 | 0.45 | 0.00 | 0.55 |
| Area STS2 (STS) right | 56 | 64 | 0.43 | 0.21 | 0.43 | 0.00 | 0.57 |
| Area Ig2 (Insula) right | 61 | 65 | 0.42 | 0.21 | 0.42 | 0.00 | 0.58 |
| Area FG1 (FusG) right | 56 | 65 | 0.40 | 0.20 | 0.40 | 0.00 | 0.60 |
| Area Id5 (Insula) right | 54 | 65 | 0.39 | 0.20 | 0.39 | 0.00 | 0.61 |
| Area hOc3v (LingG) right | 59 | 60 | 0.39 | 0.19 | 0.39 | 0.00 | 0.61 |
| Area Id4 (Insula) right | 59 | 63 | 0.38 | 0.19 | 0.38 | 0.00 | 0.62 |
| Area Fo4 (OFC) left | 55 | 65 | 0.38 | 0.19 | 0.38 | 0.00 | 0.62 |
| Area Op8 (Frontal Operculum) right | 58 | 65 | 0.38 | 0.19 | 0.38 | 0.00 | 0.62 |
| Area OP3 (POperc) right | 58 | 69 | 0.38 | 0.19 | 0.38 | 0.00 | 0.62 |
| Area 45 (IFG) right | 59 | 68 | 0.37 | 0.19 | 0.37 | 0.00 | 0.63 |
| Area OP2 (POperc) left | 58 | 67 | 0.37 | 0.19 | 0.37 | 0.00 | 0.63 |
| Frontal to Temporal I (GapMap)<br>right | 57 | 65 | 0.35 | 0.17 | 0.35 | 0.00 | 0.65 |
| Area 7A (SPL) left | 60 | 65 | 0.34 | 0.17 | 0.34 | 0.00 | 0.66 |
| Area PFt (IPL) right | 59 | 69 | 0.34 | 0.17 | 0.34 | 0.00 | 0.66 |
| Area OP4 (POperc) right | 59 | 68 | 0.34 | 0.17 | 0.34 | 0.00 | 0.66 |
| Area 44 (IFG) right | 60 | 66 | 0.34 | 0.17 | 0.34 | 0.00 | 0.66 |
| Area Id6 (Insula) left | 58 | 59 | 0.33 | 0.17 | 0.33 | 0.00 | 0.67 |
| Area Fp1 (FPole) right | 59 | 63 | 0.33 | 0.17 | 0.33 | 0.00 | 0.67 |
| HC Prosubiculum (Hippocampus)<br>right | 58 | 67 | 0.32 | 0.16 | 0.32 | 0.00 | 0.68 |
| Area OP3 (POperc) left | 54 | 65 | 0.32 | 0.16 | 0.32 | 0.00 | 0.68 |
| **Area s24 (sACC) right** | 47 | 57 | 0.32 | 0.16 | 0.32 | 0.00 | 0.68 |
| Area STS1 (STS) right | 57 | 62 | 0.32 | 0.16 | 0.32 | 0.00 | 0.68 |
| Area OP2 (POperc) right | 59 | 66 | 0.30 | 0.15 | 0.30 | 0.00 | 0.70 |
| Area hOc3d (Cuneus) right | 58 | 63 | 0.30 | 0.15 | 0.30 | 0.00 | 0.70 |
| **Area 33 (ACC) right** | 59 | 65 | 0.29 | 0.15 | 0.29 | 0.00 | 0.71 |
| Frontal to Temporal II (GapMap)<br>right | 57 | 64 | 0.29 | 0.15 | 0.29 | 0.00 | 0.71 |
| Area FG4 (FusG) left | 59 | 66 | 0.29 | 0.15 | 0.29 | 0.00 | 0.71 |
| Area PF (IPL) right | 59 | 65 | 0.29 | 0.15 | 0.29 | 0.00 | 0.71 |
| **Fastigial Nucleus (Cerebellum) left** | 56 | 62 | 0.29 | 0.14 | 0.29 | 0.00 | 0.71 |
| Area Id10 (Insula) right | 59 | 64 | 0.28 | 0.14 | 0.28 | 0.00 | 0.72 |
| Area hOc4la (LOC) left | 59 | 67 | 0.28 | 0.14 | 0.28 | 0.00 | 0.72 |
| Area Fo6 (OFC) right | 55 | 66 | 0.28 | 0.14 | 0.28 | 0.00 | 0.72 |
| Area 6d1 (PreCG) right | 57 | 64 | 0.28 | 0.14 | 0.28 | 0.00 | 0.72 |
| Area Ph1 (PhG) right | 59 | 61 | 0.27 | 0.14 | 0.27 | 0.00 | 0.73 |
| Area FG3 (FusG) left | 58 | 66 | 0.27 | 0.14 | 0.27 | 0.00 | 0.73 |
| Area 6mp (SMA mesial SFG) right | 57 | 62 | 0.26 | 0.13 | 0.26 | 0.00 | 0.74 |
| CA2 (Hippocampus) left | 59 | 67 | 0.25 | 0.13 | 0.25 | 0.00 | 0.75 |
| Area Fo6 (OFC) left | 56 | 60 | 0.24 | 0.12 | 0.24 | 0.00 | 0.76 |
| Area hIP7 (IPS) right | 61 | 62 | 0.23 | 0.12 | 0.23 | 0.00 | 0.77 |

### Sensitivity analysis

To assess the sensitivity of additive heritability in distinct regions across the brain, the data was divided into the placebo analgesia circuitry versus the rest of the brain (Figure 2). Our findings indicate that the mean heritability in placebo regions was notably higher compared to other brain areas.

## Discussion

This study is the first to directly examine the heritability of neural responses within the placebo analgesia brain circuitry, using a twin fMRI design. By quantifying genetic influences on key nodes of the placebo network, our findings provide novel evidence that individual differences in placebo analgesia are partly shaped by inherited factors.

Crucially, the ACE modeling demonstrated significant heritability at moderate levels (A = 0.21–0.49) for neural responses in an opioid-rich network underlying placebo analgesia, i.e., the rACC, pons, and cerebellar nuclei. Although placebo effects are often described as being primarily shaped by contextual factors ^15^ our study is the first twin-design to show that genetic factors also make a substantial, though not exclusive, contribution to the neural responses in brain circuitry underlying placebo analgesia. A handful of studies have examined whether variation in single-nucleotide polymorphisms (SNPs) could be associated with endogenous pain modulation ^16^ and placebo analgesia ^17^. However, SNP studies have failed to find replicable associations between genetic polymorphisms and placebo responding. Instead, it has been suggested that placebo analgesia is likely influenced by a polygenic architecture, in which multiple genes and pathways contribute to individual differences ^17^. Here, we provide evidence for a heritable component of neural activations in placebo analgesia circuitry, yet the specific genes involved in shaping these responses, and a screening panel for prediction of placebo responses, remain to be determined.

Identifying a reliable “placebo responder profile” has long been considered a central challenge of placebo research ^18^. Despite decades of work, no stable or replicable set of psychological, behavioral, or neurobiological markers has succeeded in distinguishing placebo responders from non-responders ^19^ and there are suggestions that placebo analgesia is likely to be understood as a state rather than a heritable trait ^20^. This lack of consistency highlights the complexity of expectancy-driven analgesia and suggests that key determinants of placebo responsiveness remain undiscovered. Our findings contribute a crucial piece to this puzzle by demonstrating that neural responses within the placebo analgesia circuit are shaped by additive genetic factors. By revealing heritable influences on activity in the rACC, pons, and cerebellar nuclei, this study provides evidence that individual differences in placebo responsiveness may be rooted, in part, in inherited neural profiles.

Instead of searching for a genetic profile that predicts placebo responding in general, the present experiment used a twin-design and a directed search for additive heritability in a well-established translational placebo analgesia brain network. Using this approach, we provide a new framework for understanding why some individuals benefit from expectancy-based pain modulation, such as placebo analgesia, while others do not, and open the possibility for future responder profiles that incorporate genetic and neurobiological markers alongside psychological and contextual factors^17^. Although gene–drug interactions in analgesic treatments are now well studied, emerging evidence on genetic contributions to the neural circuitry underlying placebo analgesia challenges the traditional assumption that drug and placebo responses are fundamentally distinct; one purely biological and the other purely psychological ^21^.

Our whole brain analysis, as detailed in Table 2, identified significant activation across several key brain regions during painful stimulation. Notably, multiple subregions of the anterior cingulate cortex (ACC), including right and left Area 33, p32, p24c, and p24ab, as well as the rACC exhibited robust activation. In addition, cerebellar nuclei such as the interposed, dentate, and fastigial nuclei were consistently engaged. These results underscore the central involvement of both ACC subregions and cerebellar structures in the processing and modulation of pain. The statistical values presented further support the widespread and coordinated neural response to pain within these regions, highlighting their contribution to the multifaceted experience of pain.

Our findings reveal that neural responses in the placebo analgesia circuit are shaped in part by genetic factors, adding complementary information to the recent animal work ^11,12^ and previous brain imaging studies in humans that outlined the rACC-brainstem pathways in placebo analgesia (but no previous focus on the cerebellum)^6–8^. Our results highlight the heritable contribution to an opioid-rich pain inhibitory pathway that complements our previous assessment of the heritable components of nociceptive processing in the human brain ^22^. Our results suggest that targeting the rACC–brainstem–cerebellar circuitry, conserved across species ^11,12^, may improve analgesic efficacy, particularly when interventions are tailored to genetic profiles.

### Limitations

Several limitations should be acknowledged. Although this neuroimaging twin study has a relatively large sample, increasing the cohort size would enhance statistical power to identify smaller genetic effects and address sample heterogeneity. Second, the study focused on same-sex twins aged 20–60 years, which may limit generalizability to other age groups or mixed-sex pairs. Third, the use of electrical stimulation to evoke pain during fMRI scanning, while well-controlled, may not capture the full spectrum of pain experiences encountered in clinical populations. While the study demonstrates moderate heritability within a well-documented placebo analgesia network, it is important to recognize that significant heritability was also observed in other brain regions beyond this network. This broader pattern suggests that genetic influences on pain-evoked neural responses are not exclusive to the placebo analgesic circuitry but rather extend to multiple brain systems involved in the pain experience. Future studies should aim to disentangle the unique genetic contributions to the placebo network from those affecting general pain-related brain activity, to better clarify the specificity of these findings. Additionally, it should be noted that this study was originally designed as a conditioning study with noxious stimuli, and the focus on placebo analgesia brain circuitry represents a secondary analysis. This may influence the interpretation and scope of the findings.

### Future Directions

Future research should aim to replicate these findings in larger and more diverse samples, and to integrate genetic, epigenetic, and environmental data for a more comprehensive understanding of variations in neural responses in placebo analgesia circuitry. Longitudinal studies could also elucidate how genetic and environmental influences on the placebo analgesia circuitry may predict treatment outcomes in trials where participants are treated with an active drug or placebo.

## Conclusion

In summary, this twin fMRI study demonstrates that key nodes within the placebo analgesia brain circuitry, including the rACC, pons, and cerebellar nuclei show significant, yet moderate, heritable neural responses. These findings underscore the significant role of genetic factors in shaping individual variability in placebo responsiveness. These insights have meaningful implications for the development of personalized pain management strategies, as they point toward the potential for tailoring interventions based on an individual’s genetic and neurobiological profile.

## Materials and Methods

### Subjects

Twins aged 20–60 were recruited from the Swedish Twin Registry (STR). Same-sex twin pairs with known zygosity were selected if eligible for MRI and screened for substance abuse, psychological treatment, or medications affecting emotion or cognition. After initial screening, 305 participants underwent fMRI scanning. Data were excluded due to excessive head motion (>50% frames above 0.5 mm displacement, n=16) and missing data = 3. Twins whose corresponding pair was missing were also excluded from the analysis (n = 22). The final sample included 264 twins: 62 identical pairs (35 female, 27 male; mean age 31 yeras) and 70 fraternal pairs (40 female, 30 male; mean age 31 years). All participants gave written informed consent per Uppsala Ethical Review Board guidelines and received SEK 1000 (∼100 USD) compensation. This study was pre-registered at the Open Science Foundation (OSF) (https://osf.io/hynvf/).

### Brain Imaging

Imaging data were collected with a 3.0 T Discovery MR750 scanner (GE Healthcare) and an 8-channel head coil. Foam wedges, earplugs, and headphones were used to minimize head motion and noise. T1-weighted structural images were acquired with whole head coverage (TR = 2.4 s, TE = 2.8 s, 6.04 min, flip angle 11°). Functional images used gradient EPI (TR = 2.4 s, TE = 28 ms, flip angle = 80°, 47 volumes, 3.0 mm slice thickness, axial, A/P phase-encoding). Slices were acquired interleaved and ascending. Higher order shimming and 5 dummy scans was performed before data acquisition.

### Stimuli and Contexts

Visual stimuli were displayed on a flat MR scanner screen via an Epson EX5260 projector. Stimulus presentation was run on a custom Unity 5.2.3 build and interfaced with BIOPAC for electrical stimuli through a parallel port using custom .NET serial communication software.

### fMRI Paradigm Design

Participants underwent a conditioning experiment using noxious electrical stimuli. Images of two virtual characters projected on a screen inside the scanner served as predictive cues: one as the aversive cue preceding noxious stimuli, the other as a safety cue; their assignments were counterbalanced across participants. Each cue appeared for 6 s over 16 conditioning trials, with one of the cues paired with noxious stimuli on half the presentations and not on the others. Habituation involved four presentations per cue without any noxious stimuli before conditioning. Stimulus order was varied across subjects, and each trial was followed by a jittered 8–12 s interval. The experiment lasted 9 min 47 s; habituation trials were excluded from analysis.

Noxious electrical stimuli were delivered to the distal part of the participant’s left volar forearm (adjacent to the wrist) via radio-translucent disposable dry electrodes (EL509, BIOPAC Systems, Goleta, CA). The delivery of the noxious stimuli was controlled using the STM100C module connected to the STM200 constant current stimulator (BIOPAC Systems), using a unipolar pulse with a fixed duration of 67 Hz. The voltage level was individually calibrated before the experiment using an ascending staircase procedure until stimuli were rated as “aversive” ^23^. After finding the voltage level that participants rated as aversive, this parameter was kept constant throughout the experiment. The determined average electrical voltage was *M* = 31 V, *SD* = 7, and range = (15, 55).

### Analysis of fMRI Imaging Data

fMRI data were analyzed using ENIGMA HALFpipe ^24^. Preprocessing involved slice timing correction, realignment, co-registration to T1 images, normalization to MNI152NLin6Asym space and ICA-AROMA filtering.

First-level analysis used an event-related model to estimate BOLD responses during the noxious stimulations, here referred to as pain processing. Three event types were modeled with separate regressors: the aversive cue preceding noxious stimuli, the same cue not followed by noxious stimuli and noxious stimuli. The aversive cue co-terminated with the noxious stimuli in 50% of trials; durations were 6 s for the aversive and non-aversive cue and 3 s for noxious stimuli. We assessed the additive heritability (A) by focusing on neural activations during noxious stimuli. The original aim of this experiment was to investigate fear conditioning, and not on the electrical stimuli per se. Details regarding the conditioning results can be found in Kastrati et al. ^25^. Anatomical labeling of significantly activated brain regions were performed using the Julich brain atlas v29 ^26^ and two combined masks from a probability atlas based on the Human Connectome Project ^27^. To enhance power and specificity given the small mask sizes, Brainstem pons L/R was derived from CSTL L/R and FPTL L/R masks. Supplementary Table 1 includes details on which brain regions were selected from each atlas package.

### Estimation of genetic influences on brain function

#### Outlier Identification

Prior to genetic modeling, univariate outliers were identified and removed to enhance correlation robustness between MZ and DZ twins, following recommendations for small samples ^28^ and previous neuroimaging protocols ^29^. Outliers were defined as participants with mean contrast estimates responses exceeding 1.5 times the interquartile range in any atlas mask or contrast, visualized via boxplots in R (R Core Team 2024). If one twin was an outlier, both were excluded.

#### Genetic Effects Estimation

After outlier exclusion, additive genetic effects on brain data were analyzed with the Mets package in R, decomposing variance into additive genetic, common environment, and nonshared environment/error. These factors were estimated by comparing correlations between MZ and DZ pairs, with A reflecting genetic similarity, C representing shared environment, and E unique experiences and error. Estimates were calculated for each of the 296 ROIs for the pain contrast.

## Acknowledgements

This research was supported by grants from the Swedish Research Council (2014-01160, 2018-01322) and Riksbankens Jubileumsfond (P20-0125).

## Author contributions

J.R and F.Å. designed the experiment. G.K. and J.R. performed the experiments. J.R and K.J. wrote the first draft. All authors substantially revised the manuscript. J.R analyzed the data K.J. contributed to interpretation of results. All authors have read and approved the manuscript.

## Data and materials availability

All data generated during the current study can be made available upon reasonable request from the corresponding author.

## Ethics statement

The study was approved by the Uppsala Ethical Review Board (Dnr 2014-01160).

