## Supplemental table and figure for "Genetic Influences on Neural Responses in Placebo Analgesia Circuitry"

**Supplementary materials**

**Supplemental Table 1**. Additive heritability (A), common environment (C) and error variance (E) for all brain regions included in the Julich brain atlas. (*) indicates brainstem regions identified using a probability atlas derived from the Human Connectome Project. All p-values are Bonferroni corrected ($p < 0.05$).

| Brain region | MZ-pairs | DZ-pairs | Mets MZ corr | Mets DZ corr | A | A p-values | C | E |
|---|---|---|---|---|---|---|---|---|
| Area 1 (PostCG) left | 54 | 64 | 0.14 | 0.14 | 0.00 | 1.00 | 0.14 | 0.86 |
| Area 1 (PostCG) right | 60 | 66 | 0.39 | 0.32 | 0.14 | 1.00 | 0.25 | 0.61 |
| Area 25 (sACC) left | 49 | 54 | 0.13 | 0.13 | 0.00 | 1.00 | 0.13 | 0.87 |
| Area 25 (sACC) right | 55 | 61 | 0.18 | 0.18 | 0.00 | 1.00 | 0.18 | 0.82 |
| Area 2 (PostCS) left | 57 | 66 | 0.10 | 0.05 | 0.10 | 1.00 | 0.00 | 0.90 |
| Area 2 (PostCS) right | 61 | 67 | 0.32 | 0.17 | 0.30 | 1.00 | 0.02 | 0.68 |
| Area 33 (ACC) left | 59 | 67 | 0.21 | 0.10 | 0.21 | 0.70 | 0.00 | 0.79 |
| Area 33 (ACC) right | 59 | 65 | 0.29 | 0.15 | 0.29 | 0.00 | 0.00 | 0.71 |
| Area 3a (PostCG) left | 57 | 64 | 0.05 | 0.04 | 0.03 | 1.00 | 0.02 | 0.95 |
| Area 3a (PostCG) right | 57 | 63 | 0.15 | 0.15 | 0.00 | 1.00 | 0.15 | 0.85 |
| Area 3b (PostCG) left | 58 | 67 | 0.12 | 0.06 | 0.12 | 1.00 | 0.00 | 0.88 |
| Area 3b (PostCG) right | 57 | 65 | 0.15 | 0.15 | 0.00 | 1.00 | 0.15 | 0.85 |
| Area 44 (IFG) left | 59 | 65 | 0.36 | 0.23 | 0.26 | 1.00 | 0.10 | 0.64 |
| Area 44 (IFG) right | 60 | 66 | 0.34 | 0.17 | 0.34 | 0.00 | 0.00 | 0.66 |
| Area 45 (IFG) left | 60 | 66 | 0.18 | 0.13 | 0.10 | 1.00 | 0.08 | 0.82 |
| Area 45 (IFG) right | 59 | 68 | 0.37 | 0.19 | 0.37 | 0.00 | 0.00 | 0.63 |
| Area 4a (PreCG) left | 57 | 61 | 0.08 | 0.04 | 0.08 | 1.00 | 0.00 | 0.92 |
| Area 4a (PreCG) right | 55 | 63 | 0.21 | 0.11 | 0.21 | 0.89 | 0.00 | 0.79 |
| Area 4p (PreCG) left | 55 | 63 | 0.15 | 0.09 | 0.13 | 1.00 | 0.02 | 0.85 |

| | | | | | | | | |
|---|---|---|---|---|---|---|---|---|
| Area 4p (PreCG) right | 53 | 62 | 0.33 | 0.33 | 0.00 | 1.00 | 0.33 | 0.67 |
| Area 5Ci (SPL) left | 55 | 65 | 0.11 | 0.09 | 0.03 | 1.00 | 0.08 | 0.89 |
| Area 5Ci (SPL) right | 58 | 65 | 0.12 | 0.06 | 0.12 | 1.00 | 0.00 | 0.88 |
| Area 5L (SPL) left | 56 | 64 | 0.18 | 0.09 | 0.18 | 1.00 | 0.00 | 0.82 |
| Area 5L (SPL) right | 58 | 69 | 0.24 | 0.12 | 0.23 | 1.00 | 0.00 | 0.76 |
| Area 5M (SPL) left | 55 | 65 | 0.02 | 0.02 | 0.00 | 1.00 | 0.02 | 0.98 |
| Area 5M (SPL) right | 58 | 65 | 0.11 | 0.11 | 0.00 | 1.00 | 0.11 | 0.89 |
| Area 6d1 (PreCG) left | 57 | 62 | 0.18 | 0.09 | 0.18 | 1.00 | 0.00 | 0.82 |
| Area 6d1 (PreCG) right | 57 | 64 | 0.28 | 0.14 | 0.28 | 0.03 | 0.00 | 0.72 |
| Area 6d2 (PreCG) left | 53 | 67 | 0.16 | 0.08 | 0.16 | 1.00 | 0.00 | 0.84 |
| Area 6d2 (PreCG) right | 59 | 66 | 0.15 | 0.07 | 0.15 | 1.00 | 0.00 | 0.85 |
| Area 6d3 (SFS) left | 60 | 65 | 0.09 | 0.06 | 0.06 | 1.00 | 0.03 | 0.91 |
| Area 6d3 (SFS) right | 57 | 61 | 0.22 | 0.11 | 0.22 | 0.06 | 0.00 | 0.78 |
| Area 6ma (preSMA mesial SFG) left | 58 | 67 | 0.05 | 0.03 | 0.05 | 1.00 | 0.00 | 0.95 |
| Area 6ma (preSMA mesial SFG) right | 58 | 66 | 0.18 | 0.09 | 0.18 | 1.00 | 0.00 | 0.82 |
| Area 6mp (SMA mesial SFG) left | 59 | 65 | 0.14 | 0.07 | 0.14 | 1.00 | 0.00 | 0.86 |
| Area 6mp (SMA mesial SFG) right | 57 | 62 | 0.26 | 0.13 | 0.26 | 0.00 | 0.00 | 0.74 |
| Area 7A (SPL) left | 60 | 65 | 0.34 | 0.17 | 0.34 | 0.00 | 0.00 | 0.66 |
| Area 7A (SPL) right | 60 | 68 | 0.23 | 0.22 | 0.01 | 1.00 | 0.22 | 0.77 |
| Area 7M (SPL) left | 55 | 62 | 0.12 | 0.06 | 0.12 | 1.00 | 0.00 | 0.88 |
| Area 7M (SPL) right | 58 | 63 | 0.15 | 0.07 | 0.15 | 1.00 | 0.00 | 0.85 |
| Area 7PC (SPL) left | 59 | 64 | 0.14 | 0.09 | 0.11 | 1.00 | 0.03 | 0.86 |
| Area 7PC (SPL) right | 59 | 66 | 0.14 | 0.14 | 0.00 | 1.00 | 0.14 | 0.86 |
| Area 7P (SPL) left | 57 | 64 | 0.15 | 0.15 | 0.00 | 1.00 | 0.15 | 0.85 |
| Area 7P (SPL) right | 57 | 67 | 0.11 | 0.11 | 0.00 | 1.00 | 0.11 | 0.89 |
| Area 8d1 (SFG) left | 60 | 67 | 0.19 | 0.10 | 0.19 | 1.00 | 0.00 | 0.81 |
| Area 8d1 (SFG) right | 56 | 63 | 0.00 | 0.00 | 0.00 | 1.00 | 0.00 | 1.00 |
| Area 8d2 (SFG) left | 61 | 62 | 0.05 | 0.05 | 0.00 | 1.00 | 0.05 | 0.95 |

| | | | | | | | | |
|---|---|---|---|---|---|---|---|---|
| Area 8d2 (SFG) right | 60 | 61 | 0.04 | 0.02 | 0.04 | 1.00 | 0.00 | 0.96 |
| Area 8v1 (MFG) left | 60 | 63 | 0.00 | 0.00 | 0.00 | 1.00 | 0.00 | 1.00 |
| Area 8v1 (MFG) right | 58 | 63 | 0.00 | 0.00 | 0.00 | 1.00 | 0.00 | 1.00 |
| Area 8v2 (MFG) left | 59 | 66 | 0.01 | 0.01 | 0.00 | 1.00 | 0.01 | 0.99 |
| Area 8v2 (MFG) right | 57 | 60 | 0.02 | 0.01 | 0.02 | 1.00 | 0.00 | 0.98 |
| Area CoS1 (CoS) left | 59 | 55 | 0.01 | 0.01 | 0.00 | 1.00 | 0.01 | 0.99 |
| Area CoS1 (CoS) right | 49 | 54 | 0.00 | 0.00 | 0.00 | 1.00 | 0.00 | 1.00 |
| Area FG1 (FusG) left | 59 | 64 | 0.16 | 0.08 | 0.16 | 1.00 | 0.00 | 0.84 |
| Area FG1 (FusG) right | 56 | 65 | 0.40 | 0.20 | 0.40 | 0.00 | 0.00 | 0.60 |
| Area FG2 (FusG) left | 58 | 67 | 0.18 | 0.09 | 0.18 | 1.00 | 0.00 | 0.82 |
| Area FG2 (FusG) right | 61 | 64 | 0.04 | 0.02 | 0.04 | 1.00 | 0.00 | 0.96 |
| Area FG3 (FusG) left | 58 | 66 | 0.27 | 0.14 | 0.27 | 0.00 | 0.00 | 0.73 |
| Area FG3 (FusG) right | 60 | 61 | 0.17 | 0.09 | 0.17 | 0.29 | 0.00 | 0.83 |
| Area FG4 (FusG) left | 59 | 66 | 0.29 | 0.15 | 0.29 | 0.01 | 0.00 | 0.71 |
| Area FG4 (FusG) right | 60 | 65 | 0.13 | 0.07 | 0.13 | 1.00 | 0.00 | 0.87 |
| Area Fo1 (OFC) left | 44 | 47 | 0.00 | 0.00 | 0.00 | 1.00 | 0.00 | 1.00 |
| Area Fo1 (OFC) right | 23 | 32 | 0.00 | 0.00 | 0.00 | 1.00 | 0.00 | 1.00 |
| Area Fo2 (OFC) left | 11 | 12 | 0.11 | 0.05 | 0.11 | 1.00 | 0.00 | 0.89 |
| Area Fo2 (OFC) right | 13 | 19 | 0.02 | 0.02 | 0.00 | 1.00 | 0.02 | 0.98 |
| Area Fo3 (OFC) left | 58 | 66 | 0.07 | 0.03 | 0.07 | 1.00 | 0.00 | 0.93 |
| Area Fo3 (OFC) right | 56 | 61 | 0.23 | 0.11 | 0.23 | 0.10 | 0.00 | 0.77 |
| Area Fo4 (OFC) left | 55 | 65 | 0.38 | 0.19 | 0.38 | 0.00 | 0.00 | 0.62 |
| Area Fo4 (OFC) right | 55 | 63 | 0.04 | 0.02 | 0.04 | 1.00 | 0.00 | 0.96 |
| Area Fo5 (OFC) left | 55 | 67 | 0.00 | 0.00 | 0.00 | 1.00 | 0.00 | 1.00 |
| Area Fo5 (OFC) right | 58 | 65 | 0.19 | 0.09 | 0.19 | 1.00 | 0.00 | 0.81 |
| Area Fo6 (OFC) left | 56 | 60 | 0.24 | 0.12 | 0.24 | 0.03 | 0.00 | 0.76 |
| Area Fo6 (OFC) right | 55 | 66 | 0.28 | 0.14 | 0.28 | 0.03 | 0.00 | 0.72 |
| Area Fo7 (OFC) left | 55 | 66 | 0.23 | 0.16 | 0.15 | 1.00 | 0.08 | 0.77 |

| | | | | | | | | |
|---|---|---|---|---|---|---|---|---|
| Area Fo7 (OFC) right | 57 | 62 | 0.23 | 0.11 | 0.23 | 0.07 | 0.00 | 0.77 |
| Area Fp1 (FPole) left | 56 | 61 | 0.26 | 0.13 | 0.26 | 0.11 | 0.00 | 0.74 |
| Area Fp1 (FPole) right | 59 | 63 | 0.33 | 0.17 | 0.33 | 0.01 | 0.00 | 0.67 |
| Area Fp2 (FPole) left | 59 | 63 | 0.21 | 0.10 | 0.21 | 0.42 | 0.00 | 0.79 |
| Area Fp2 (FPole) right | 61 | 58 | 0.21 | 0.12 | 0.19 | 1.00 | 0.02 | 0.79 |
| Area IFJ1 (IFS PreCS) left | 61 | 60 | 0.15 | 0.08 | 0.15 | 1.00 | 0.00 | 0.85 |
| Area IFJ1 (IFS PreCS) right | 61 | 68 | 0.00 | 0.00 | 0.00 | 1.00 | 0.00 | 1.00 |
| Area IFJ2 (IFS PreCS) left | 59 | 66 | 0.49 | 0.24 | 0.49 | 0.00 | 0.00 | 0.51 |
| Area IFJ2 (IFS PreCS) right | 56 | 65 | 0.06 | 0.03 | 0.06 | 1.00 | 0.00 | 0.94 |
| Area IFS1 (IFS) left | 56 | 66 | 0.09 | 0.05 | 0.09 | 1.00 | 0.00 | 0.91 |
| Area IFS1 (IFS) right | 58 | 65 | 0.00 | 0.00 | 0.00 | 1.00 | 0.00 | 1.00 |
| Area IFS2 (IFS) left | 56 | 64 | 0.22 | 0.12 | 0.22 | 1.00 | 0.01 | 0.78 |
| Area IFS2 (IFS) right | 54 | 67 | 0.00 | 0.00 | 0.00 | 1.00 | 0.00 | 1.00 |
| Area IFS3 (IFS) left | 57 | 64 | 0.01 | 0.01 | 0.01 | 1.00 | 0.00 | 0.99 |
| Area IFS3 (IFS) right | 58 | 67 | 0.10 | 0.05 | 0.10 | 1.00 | 0.00 | 0.90 |
| Area IFS4 (IFS) left | 58 | 64 | 0.04 | 0.04 | 0.00 | 1.00 | 0.04 | 0.96 |
| Area IFS4 (IFS) right | 60 | 65 | 0.06 | 0.03 | 0.06 | 1.00 | 0.00 | 0.94 |
| Area Ia1 (Insula) left | 58 | 68 | 0.09 | 0.09 | 0.00 | 1.00 | 0.09 | 0.91 |
| Area Ia1 (Insula) right | 56 | 63 | 0.16 | 0.08 | 0.16 | 1.00 | 0.00 | 0.84 |
| Area Ia2 (Insula) left | 54 | 63 | 0.05 | 0.02 | 0.05 | 1.00 | 0.00 | 0.95 |
| Area Ia2 (Insula) right | 61 | 66 | 0.11 | 0.05 | 0.11 | 1.00 | 0.00 | 0.89 |
| Area Ia3 (Insula) left | 55 | 65 | 0.17 | 0.09 | 0.17 | 1.00 | 0.00 | 0.83 |
| Area Ia3 (Insula) right | 56 | 69 | 0.11 | 0.06 | 0.11 | 1.00 | 0.00 | 0.89 |
| Area Id10 (Insula) left | 56 | 61 | 0.09 | 0.05 | 0.09 | 1.00 | 0.00 | 0.91 |
| Area Id10 (Insula) right | 59 | 64 | 0.28 | 0.14 | 0.28 | 0.00 | 0.00 | 0.72 |
| Area Id1 (Insula) left | 58 | 64 | 0.25 | 0.12 | 0.25 | 0.05 | 0.00 | 0.75 |
| Area Id1 (Insula) right | 62 | 65 | 0.15 | 0.07 | 0.15 | 1.00 | 0.00 | 0.85 |
| Area Id2 (Insula) left | 60 | 68 | 0.04 | 0.02 | 0.04 | 1.00 | 0.00 | 0.96 |

| | | | | | | | | |
|---|---|---|---|---|---|---|---|---|
| Area Id2 (Insula) right | 57 | 64 | 0.17 | 0.08 | 0.17 | 1.00 | 0.00 | 0.83 |
| Area Id3 (Insula) left | 59 | 65 | 0.18 | 0.18 | 0.00 | 1.00 | 0.18 | 0.82 |
| Area Id3 (Insula) right | 58 | 63 | 0.21 | 0.11 | 0.21 | 0.34 | 0.00 | 0.79 |
| Area Id4 (Insula) left | 59 | 55 | 0.21 | 0.11 | 0.21 | 0.06 | 0.00 | 0.79 |
| Area Id4 (Insula) right | 59 | 63 | 0.38 | 0.19 | 0.38 | 0.00 | 0.00 | 0.62 |
| Area Id5 (Insula) left | 57 | 64 | 0.15 | 0.15 | 0.00 | 1.00 | 0.15 | 0.85 |
| Area Id5 (Insula) right | 54 | 65 | 0.39 | 0.20 | 0.39 | 0.00 | 0.00 | 0.61 |
| Area Id6 (Insula) left | 58 | 59 | 0.33 | 0.17 | 0.33 | 0.00 | 0.00 | 0.67 |
| Area Id6 (Insula) right | 58 | 64 | 0.03 | 0.02 | 0.03 | 1.00 | 0.00 | 0.97 |
| Area Id7 (Insula) left | 59 | 64 | 0.16 | 0.08 | 0.16 | 1.00 | 0.00 | 0.84 |
| Area Id7 (Insula) right | 58 | 68 | 0.12 | 0.06 | 0.12 | 1.00 | 0.00 | 0.88 |
| Area Id8 (Insula) left | 59 | 63 | 0.06 | 0.03 | 0.06 | 1.00 | 0.00 | 0.94 |
| Area Id8 (Insula) right | 58 | 61 | 0.08 | 0.04 | 0.08 | 1.00 | 0.00 | 0.92 |
| Area Id9 (Insula) left | 58 | 64 | 0.03 | 0.03 | 0.00 | 1.00 | 0.03 | 0.97 |
| Area Id9 (Insula) right | 57 | 65 | 0.24 | 0.12 | 0.24 | 0.26 | 0.00 | 0.76 |
| Area Ig1 (Insula) left | 59 | 66 | 0.21 | 0.10 | 0.21 | 0.07 | 0.00 | 0.79 |
| Area Ig1 (Insula) right | 60 | 62 | 0.20 | 0.10 | 0.20 | 0.17 | 0.00 | 0.80 |
| Area Ig2 (Insula) left | 60 | 65 | 0.21 | 0.11 | 0.21 | 0.24 | 0.00 | 0.79 |
| Area Ig2 (Insula) right | 61 | 65 | 0.42 | 0.21 | 0.42 | 0.00 | 0.00 | 0.58 |
| Area Ig3 (Insula) left | 61 | 63 | 0.10 | 0.10 | 0.00 | 1.00 | 0.10 | 0.90 |
| Area Ig3 (Insula) right | 57 | 65 | 0.05 | 0.02 | 0.05 | 1.00 | 0.00 | 0.95 |
| Area OP1 (POperc) left | 59 | 61 | 0.29 | 0.17 | 0.23 | 1.00 | 0.05 | 0.71 |
| Area OP1 (POperc) right | 57 | 64 | 0.22 | 0.11 | 0.22 | 0.09 | 0.00 | 0.78 |
| Area OP2 (POperc) left | 58 | 67 | 0.37 | 0.19 | 0.37 | 0.00 | 0.00 | 0.63 |
| Area OP2 (POperc) right | 59 | 66 | 0.30 | 0.15 | 0.30 | 0.00 | 0.00 | 0.70 |
| Area OP3 (POperc) left | 54 | 65 | 0.32 | 0.16 | 0.32 | 0.00 | 0.00 | 0.68 |
| Area OP3 (POperc) right | 58 | 69 | 0.38 | 0.19 | 0.38 | 0.00 | 0.00 | 0.62 |
| Area OP4 (POperc) left | 59 | 62 | 0.18 | 0.18 | 0.00 | 1.00 | 0.18 | 0.82 |

| | | | | | | | | |
|---|---|---|---|---|---|---|---|---|
| Area OP4 (POperc) right | 59 | 68 | 0.34 | 0.17 | 0.34 | 0.00 | 0.00 | 0.66 |
| Area Op5 (Frontal Operculum) left | 58 | 63 | 0.11 | 0.05 | 0.11 | 1.00 | 0.00 | 0.89 |
| Area Op5 (Frontal Operculum) right | 59 | 67 | 0.11 | 0.05 | 0.11 | 1.00 | 0.00 | 0.89 |
| Area Op6 (Frontal Operculum) left | 59 | 63 | 0.17 | 0.08 | 0.17 | 1.00 | 0.00 | 0.83 |
| Area Op6 (Frontal Operculum) right | 61 | 66 | 0.22 | 0.11 | 0.22 | 0.11 | 0.00 | 0.78 |
| Area Op7 (Frontal Operculum) left | 60 | 66 | 0.24 | 0.18 | 0.11 | 1.00 | 0.13 | 0.76 |
| Area Op7 (Frontal Operculum) right | 60 | 65 | 0.05 | 0.05 | 0.00 | 1.00 | 0.05 | 0.95 |
| Area Op8 (Frontal Operculum) left | 55 | 61 | 0.12 | 0.06 | 0.12 | 1.00 | 0.00 | 0.88 |
| Area Op8 (Frontal Operculum) right | 58 | 65 | 0.38 | 0.19 | 0.38 | 0.00 | 0.00 | 0.62 |
| Area Op9 (Frontal Operculum) left | 57 | 66 | 0.12 | 0.06 | 0.12 | 1.00 | 0.00 | 0.88 |
| Area Op9 (Frontal Operculum) right | 54 | 65 | 0.19 | 0.09 | 0.19 | 1.00 | 0.00 | 0.81 |
| Area PF (IPL) left | 59 | 61 | 0.10 | 0.05 | 0.10 | 1.00 | 0.00 | 0.90 |
| Area PF (IPL) right | 59 | 65 | 0.29 | 0.15 | 0.29 | 0.00 | 0.00 | 0.71 |
| Area PFcm (IPL) left | 59 | 66 | 0.04 | 0.02 | 0.04 | 1.00 | 0.00 | 0.96 |
| Area PFcm (IPL) right | 61 | 63 | 0.25 | 0.15 | 0.20 | 1.00 | 0.06 | 0.75 |
| Area PFm (IPL) left | 57 | 64 | 0.09 | 0.04 | 0.09 | 1.00 | 0.00 | 0.91 |
| Area PFm (IPL) right | 61 | 66 | 0.02 | 0.01 | 0.02 | 1.00 | 0.00 | 0.98 |
| Area PFop (IPL) left | 58 | 67 | 0.18 | 0.18 | 0.00 | 1.00 | 0.18 | 0.82 |
| Area PFop (IPL) right | 59 | 68 | 0.20 | 0.10 | 0.20 | 0.52 | 0.00 | 0.80 |
| Area PFt (IPL) left | 59 | 68 | 0.07 | 0.04 | 0.06 | 1.00 | 0.00 | 0.93 |
| Area PFt (IPL) right | 59 | 69 | 0.34 | 0.17 | 0.34 | 0.00 | 0.00 | 0.66 |
| Area PGa (IPL) left | 58 | 65 | 0.09 | 0.04 | 0.09 | 1.00 | 0.00 | 0.91 |
| Area PGa (IPL) right | 60 | 65 | 0.07 | 0.04 | 0.07 | 1.00 | 0.00 | 0.93 |
| Area PGp (IPL) left | 59 | 67 | 0.07 | 0.03 | 0.07 | 1.00 | 0.00 | 0.93 |
| Area PGp (IPL) right | 55 | 65 | 0.25 | 0.12 | 0.25 | 0.31 | 0.00 | 0.75 |
| Area Ph1 (PhG) left | 59 | 63 | 0.11 | 0.05 | 0.11 | 1.00 | 0.00 | 0.89 |
| Area Ph1 (PhG) right | 59 | 61 | 0.27 | 0.14 | 0.27 | 0.00 | 0.00 | 0.73 |
| Area Ph2 (PhG) left | 52 | 68 | 0.00 | 0.00 | 0.00 | 1.00 | 0.00 | 1.00 |

| | | | | | | | | |
|---|---|---|---|---|---|---|---|---|
| Area Ph2 (PhG) right | 59 | 63 | 0.27 | 0.19 | 0.17 | 1.00 | 0.10 | 0.73 |
| Area Ph3 (PhG) left | 53 | 66 | 0.09 | 0.05 | 0.09 | 1.00 | 0.00 | 0.91 |
| Area Ph3 (PhG) right | 59 | 64 | 0.11 | 0.06 | 0.11 | 1.00 | 0.00 | 0.89 |
| Area STS1 (STS) left | 56 | 65 | 0.11 | 0.11 | 0.00 | 1.00 | 0.11 | 0.89 |
| Area STS1 (STS) right | 57 | 62 | 0.32 | 0.16 | 0.32 | 0.00 | 0.00 | 0.68 |
| Area STS2 (STS) left | 61 | 64 | 0.09 | 0.09 | 0.00 | 1.00 | 0.09 | 0.91 |
| Area STS2 (STS) right | 56 | 64 | 0.43 | 0.21 | 0.43 | 0.00 | 0.00 | 0.57 |
| Area TE 1 0 (HESCHL) left | 58 | 64 | 0.12 | 0.12 | 0.00 | 1.00 | 0.12 | 0.88 |
| Area TE 1 0 (HESCHL) right | 59 | 63 | 0.14 | 0.10 | 0.07 | 1.00 | 0.07 | 0.86 |
| Area TE 1 1 (HESCHL) left | 52 | 62 | 0.00 | 0.00 | 0.00 | 1.00 | 0.00 | 1.00 |
| Area TE 1 1 (HESCHL) right | 59 | 63 | 0.00 | 0.00 | 0.00 | 1.00 | 0.00 | 1.00 |
| Area TE 1 2 (HESCHL) left | 60 | 65 | 0.22 | 0.22 | 0.00 | 1.00 | 0.22 | 0.78 |
| Area TE 1 2 (HESCHL) right | 58 | 64 | 0.04 | 0.04 | 0.00 | 1.00 | 0.04 | 0.96 |
| Area TE 2 1 (STG) left | 55 | 61 | 0.20 | 0.20 | 0.00 | 1.00 | 0.20 | 0.80 |
| Area TE 2 1 (STG) right | 59 | 64 | 0.10 | 0.05 | 0.10 | 1.00 | 0.00 | 0.90 |
| Area TE 2 2 (STG) left | 58 | 64 | 0.16 | 0.08 | 0.16 | 1.00 | 0.00 | 0.84 |
| Area TE 2 2 (STG) right | 58 | 63 | 0.25 | 0.12 | 0.25 | 0.45 | 0.00 | 0.75 |
| Area TE 3 (STG) left | 58 | 64 | 0.21 | 0.10 | 0.21 | 0.25 | 0.00 | 0.79 |
| Area TE 3 (STG) right | 56 | 66 | 0.03 | 0.03 | 0.00 | 1.00 | 0.03 | 0.97 |
| Area TI (STG) left | 56 | 62 | 0.00 | 0.00 | 0.00 | 1.00 | 0.00 | 1.00 |
| Area TI (STG) right | 59 | 57 | 0.00 | 0.00 | 0.00 | 1.00 | 0.00 | 1.00 |
| Area TPJ (STG SMG) left | 60 | 64 | 0.00 | 0.00 | 0.00 | 1.00 | 0.00 | 1.00 |
| Area TPJ (STG SMG) right | 57 | 60 | 0.01 | 0.01 | 0.01 | 1.00 | 0.00 | 0.99 |
| Area TeI (STG) left | 58 | 61 | 0.00 | 0.00 | 0.00 | 1.00 | 0.00 | 1.00 |
| Area TeI (STG) right | 55 | 65 | 0.20 | 0.10 | 0.20 | 1.00 | 0.00 | 0.80 |
| Area hIP1 (IPS) left | 59 | 65 | 0.03 | 0.03 | 0.00 | 1.00 | 0.03 | 0.97 |
| Area hIP1 (IPS) right | 57 | 65 | 0.00 | 0.00 | 0.00 | 1.00 | 0.00 | 1.00 |
| Area hIP2 (IPS) left | 58 | 65 | 0.00 | 0.00 | 0.00 | 1.00 | 0.00 | 1.00 |

| | | | | | | | | |
|---|---|---|---|---|---|---|---|---|
| Area hIP2 (IPS) right | 56 | 67 | 0.00 | 0.00 | 0.00 | 1.00 | 0.00 | 1.00 |
| Area hIP3 (IPS) left | 61 | 60 | 0.03 | 0.02 | 0.01 | 1.00 | 0.02 | 0.97 |
| Area hIP3 (IPS) right | 60 | 63 | 0.07 | 0.04 | 0.07 | 1.00 | 0.00 | 0.93 |
| Area hIP4 (IPS) left | 57 | 64 | 0.16 | 0.16 | 0.00 | 1.00 | 0.16 | 0.84 |
| Area hIP4 (IPS) right | 57 | 66 | 0.10 | 0.05 | 0.10 | 1.00 | 0.00 | 0.90 |
| Area hIP5 (IPS) left | 61 | 68 | 0.07 | 0.04 | 0.07 | 1.00 | 0.00 | 0.93 |
| Area hIP5 (IPS) right | 60 | 68 | 0.00 | 0.00 | 0.00 | 1.00 | 0.00 | 1.00 |
| Area hIP6 (IPS) left | 61 | 68 | 0.11 | 0.06 | 0.11 | 1.00 | 0.00 | 0.89 |
| Area hIP6 (IPS) right | 58 | 68 | 0.00 | 0.00 | 0.00 | 1.00 | 0.00 | 1.00 |
| Area hIP7 (IPS) left | 60 | 60 | 0.13 | 0.06 | 0.13 | 1.00 | 0.00 | 0.87 |
| Area hIP7 (IPS) right | 61 | 62 | 0.23 | 0.12 | 0.23 | 0.03 | 0.00 | 0.77 |
| Area hIP8 (IPS) left | 59 | 63 | 0.19 | 0.14 | 0.10 | 1.00 | 0.09 | 0.81 |
| Area hIP8 (IPS) right | 60 | 68 | 0.01 | 0.01 | 0.01 | 1.00 | 0.00 | 0.99 |
| Area hOc1 (V1 17 CalcS) left | 58 | 67 | 0.28 | 0.14 | 0.28 | 1.00 | 0.00 | 0.72 |
| Area hOc1 (V1 17 CalcS) right | 58 | 67 | 0.20 | 0.10 | 0.20 | 0.77 | 0.00 | 0.80 |
| Area hOc2 (V2 18) left | 60 | 66 | 0.17 | 0.08 | 0.17 | 1.00 | 0.00 | 0.83 |
| Area hOc2 (V2 18) right | 61 | 66 | 0.25 | 0.14 | 0.22 | 1.00 | 0.03 | 0.75 |
| Area hOc3d (Cuneus) left | 59 | 64 | 0.17 | 0.09 | 0.17 | 1.00 | 0.00 | 0.83 |
| Area hOc3d (Cuneus) right | 58 | 63 | 0.30 | 0.15 | 0.30 | 0.00 | 0.00 | 0.70 |
| Area hOc3v (LingG) left | 58 | 64 | 0.13 | 0.07 | 0.13 | 1.00 | 0.00 | 0.87 |
| Area hOc3v (LingG) right | 59 | 60 | 0.39 | 0.19 | 0.39 | 0.00 | 0.00 | 0.61 |
| Area hOc4d (Cuneus) left | 58 | 64 | 0.11 | 0.05 | 0.11 | 1.00 | 0.00 | 0.89 |
| Area hOc4d (Cuneus) right | 61 | 60 | 0.17 | 0.16 | 0.03 | 1.00 | 0.14 | 0.83 |
| Area hOc4la (LOC) left | 59 | 67 | 0.28 | 0.14 | 0.28 | 0.00 | 0.00 | 0.72 |
| Area hOc4la (LOC) right | 59 | 64 | 0.29 | 0.19 | 0.21 | 1.00 | 0.09 | 0.71 |
| Area hOc4lp (LOC) left | 61 | 65 | 0.13 | 0.07 | 0.13 | 1.00 | 0.00 | 0.87 |
| Area hOc4lp (LOC) right | 61 | 63 | 0.11 | 0.06 | 0.11 | 1.00 | 0.00 | 0.89 |
| Area hOc4v (LingG) left | 59 | 63 | 0.00 | 0.00 | 0.00 | 1.00 | 0.00 | 1.00 |

| | | | | | | | | |
|---|---|---|---|---|---|---|---|---|
| Area hOc4v (LingG) right | 58 | 65 | 0.00 | 0.00 | 0.00 | 1.00 | 0.00 | 1.00 |
| Area hOc5 (LOC) left | 61 | 68 | 0.17 | 0.08 | 0.17 | 1.00 | 0.00 | 0.83 |
| Area hOc5 (LOC) right | 59 | 62 | 0.22 | 0.11 | 0.22 | 0.33 | 0.00 | 0.78 |
| Area hOc6 (POS) left | 57 | 59 | 0.06 | 0.06 | 0.00 | 1.00 | 0.06 | 0.94 |
| Area hOc6 (POS) right | 58 | 67 | 0.29 | 0.28 | 0.00 | 1.00 | 0.28 | 0.71 |
| Area hPO1 (POS) left | 57 | 61 | 0.00 | 0.00 | 0.00 | 1.00 | 0.00 | 1.00 |
| Area hPO1 (POS) right | 57 | 55 | 0.17 | 0.09 | 0.17 | 1.00 | 0.00 | 0.83 |
| Area p24ab (pACC) left | 58 | 64 | 0.13 | 0.07 | 0.12 | 1.00 | 0.01 | 0.87 |
| Area p24ab (pACC) right | 58 | 64 | 0.09 | 0.09 | 0.00 | 1.00 | 0.09 | 0.91 |
| Area p24c (pACC) left | 57 | 63 | 0.12 | 0.12 | 0.00 | 1.00 | 0.12 | 0.88 |
| Area p24c (pACC) right | 59 | 64 | 0.12 | 0.06 | 0.12 | 1.00 | 0.00 | 0.88 |
| Area p32 (pACC) left | 60 | 64 | 0.27 | 0.19 | 0.16 | 1.00 | 0.11 | 0.73 |
| Area p32 (pACC) right | 61 | 66 | 0.20 | 0.10 | 0.20 | 1.00 | 0.00 | 0.80 |
| Area s24 (sACC) left | 36 | 41 | 0.10 | 0.10 | 0.00 | 1.00 | 0.10 | 0.90 |
| Area s24 (sACC) right | 47 | 57 | 0.32 | 0.16 | 0.32 | 0.00 | 0.00 | 0.68 |
| Area s32 (sACC) left | 53 | 61 | 0.21 | 0.11 | 0.21 | 1.00 | 0.00 | 0.79 |
| Area s32 (sACC) right | 51 | 54 | 0.01 | 0.01 | 0.01 | 1.00 | 0.00 | 0.99 |
| BST (Bed Nucleus) left | 55 | 63 | 0.13 | 0.06 | 0.13 | 1.00 | 0.00 | 0.87 |
| BST (Bed Nucleus) right | 58 | 68 | 0.00 | 0.00 | 0.00 | 1.00 | 0.00 | 1.00 |
| CA1 (Hippocampus) left | 60 | 65 | 0.06 | 0.03 | 0.06 | 1.00 | 0.00 | 0.94 |
| CA1 (Hippocampus) right | 55 | 67 | 0.29 | 0.15 | 0.29 | 0.09 | 0.00 | 0.71 |
| CA2 (Hippocampus) left | 59 | 67 | 0.25 | 0.13 | 0.25 | 0.00 | 0.00 | 0.75 |
| CA2 (Hippocampus) right | 58 | 67 | 0.09 | 0.05 | 0.09 | 1.00 | 0.00 | 0.91 |
| CA3 (Hippocampus) left | 59 | 66 | 0.03 | 0.02 | 0.03 | 1.00 | 0.00 | 0.97 |
| CA3 (Hippocampus) right | 59 | 66 | 0.06 | 0.03 | 0.06 | 1.00 | 0.00 | 0.94 |
| CGL (Metathalamus) left | 52 | 68 | 0.00 | 0.00 | 0.00 | 1.00 | 0.00 | 1.00 |
| CGL (Metathalamus) right | 55 | 65 | 0.11 | 0.10 | 0.01 | 1.00 | 0.09 | 0.89 |
| CGM (Metathalamus) left | 60 | 61 | 0.14 | 0.07 | 0.14 | 1.00 | 0.00 | 0.86 |

| | | | | | | | | |
|---|---|---|---|---|---|---|---|---|
| CGM (Metathalamus) right | 57 | 67 | 0.16 | 0.08 | 0.16 | 1.00 | 0.00 | 0.84 |
| CM (Amygdala) left | 8 | 10 | 0.00 | 0.00 | 0.00 | 1.00 | 0.00 | 1.00 |
| CM (Amygdala) right | 28 | 37 | 0.07 | 0.03 | 0.07 | 1.00 | 0.00 | 0.93 |
| Ch 123 (Basal Forebrain) left | 53 | 65 | 0.00 | 0.00 | 0.00 | 1.00 | 0.00 | 1.00 |
| Ch 123 (Basal Forebrain) right | 57 | 65 | 0.01 | 0.01 | 0.00 | 1.00 | 0.01 | 0.99 |
| Ch 4 (Basal Forebrain) left | 51 | 64 | 0.00 | 0.00 | 0.00 | 1.00 | 0.00 | 1.00 |
| Ch 4 (Basal Forebrain) right | 58 | 65 | 0.02 | 0.01 | 0.02 | 1.00 | 0.00 | 0.98 |
| DG (Hippocampus) left | 54 | 66 | 0.24 | 0.12 | 0.24 | 0.21 | 0.00 | 0.76 |
| DG (Hippocampus) right | 57 | 65 | 0.16 | 0.08 | 0.16 | 1.00 | 0.00 | 0.84 |
| Dorsal Dentate Nucleus (Cerebellum) left | 57 | 64 | 0.06 | 0.03 | 0.06 | 1.00 | 0.00 | 0.94 |
| Dorsal Dentate Nucleus (Cerebellum) right | 57 | 59 | 0.09 | 0.09 | 0.00 | 1.00 | 0.09 | 0.91 |
| Entorhinal Cortex left | 10 | 11 | 0.00 | 0.00 | 0.00 | 1.00 | 0.00 | 1.00 |
| Entorhinal Cortex right | 11 | 12 | 0.00 | 0.00 | 0.00 | 1.00 | 0.00 | 1.00 |
| Fastigial Nucleus (Cerebellum) left | 56 | 62 | 0.29 | 0.14 | 0.29 | 0.00 | 0.00 | 0.71 |
| Fastigial Nucleus (Cerebellum) right | 58 | 65 | 0.00 | 0.00 | 0.00 | 1.00 | 0.00 | 1.00 |
| Frontal II (GapMap) left | 60 | 66 | 0.03 | 0.03 | 0.00 | 1.00 | 0.03 | 0.97 |
| Frontal II (GapMap) right | 58 | 66 | 0.22 | 0.11 | 0.22 | 0.35 | 0.00 | 0.78 |
| Frontal I (GapMap) left | 56 | 64 | 0.30 | 0.15 | 0.30 | 0.25 | 0.00 | 0.70 |
| Frontal I (GapMap) right | 59 | 63 | 0.19 | 0.09 | 0.19 | 1.00 | 0.00 | 0.81 |
| Frontal to Occipital (GapMap) left | 56 | 65 | 0.21 | 0.11 | 0.21 | 0.58 | 0.00 | 0.79 |
| Frontal to Occipital (GapMap) right | 56 | 68 | 0.07 | 0.04 | 0.07 | 1.00 | 0.00 | 0.93 |
| Frontal to Temporal II (GapMap) left | 57 | 64 | 0.21 | 0.11 | 0.21 | 0.73 | 0.00 | 0.79 |
| Frontal to Temporal II (GapMap) right | 57 | 64 | 0.29 | 0.15 | 0.29 | 0.00 | 0.00 | 0.71 |
| Frontal to Temporal I (GapMap) left | 59 | 65 | 0.22 | 0.11 | 0.22 | 0.57 | 0.00 | 0.78 |
| Frontal to Temporal I (GapMap) right | 57 | 65 | 0.35 | 0.17 | 0.35 | 0.00 | 0.00 | 0.65 |
| HATA (Hippocampus) left | 49 | 49 | 0.00 | 0.00 | 0.00 | 1.00 | 0.00 | 1.00 |
| HATA (Hippocampus) right | 59 | 64 | 0.12 | 0.06 | 0.12 | 1.00 | 0.00 | 0.88 |
| HC Parasubiculum (Hippocampus) left | 7 | 11 | 0.59 | 0.29 | 0.59 | 1.00 | 0.00 | 0.41 |

| | | | | | | | |
|---|---|---|---|---|---|---|---|
| HC Parasubiculum (Hippocampus) right | 34 | 37 | 0.00 | 0.00 | 0.00 | 1.00 | 0.00 | 1.00 |
| HC Presubiculum (Hippocampus) left | 59 | 67 | 0.06 | 0.03 | 0.06 | 1.00 | 0.00 | 0.94 |
| HC Presubiculum (Hippocampus) right | 57 | 66 | 0.00 | 0.00 | 0.00 | 1.00 | 0.00 | 1.00 |
| HC Prosubiculum (Hippocampus) left | 50 | 61 | 0.08 | 0.08 | 0.00 | 1.00 | 0.08 | 0.92 |
| HC Prosubiculum (Hippocampus) right | 58 | 67 | 0.32 | 0.16 | 0.32 | 0.00 | 0.00 | 0.68 |
| HC Subiculum (Hippocampus) left | 56 | 67 | 0.01 | 0.00 | 0.01 | 1.00 | 0.00 | 0.99 |
| HC Subiculum (Hippocampus) right | 54 | 66 | 0.21 | 0.10 | 0.21 | 0.66 | 0.00 | 0.79 |
| HC Transsubiculum (Hippocampus) left | 59 | 66 | 0.09 | 0.09 | 0.00 | 1.00 | 0.09 | 0.91 |
| HC Transsubiculum (Hippocampus) right | 57 | 64 | 0.05 | 0.05 | 0.00 | 1.00 | 0.05 | 0.95 |
| Interposed Nucleus (Cerebellum) left | 59 | 62 | 0.00 | 0.00 | 0.00 | 1.00 | 0.00 | 1.00 |
| Interposed Nucleus (Cerebellum) right | 52 | 63 | 0.15 | 0.07 | 0.15 | 1.00 | 0.00 | 0.85 |
| LB (Amygdala) left | 22 | 25 | 0.00 | 0.00 | 0.00 | 1.00 | 0.00 | 1.00 |
| LB (Amygdala) right | 58 | 60 | 0.07 | 0.04 | 0.07 | 1.00 | 0.00 | 0.93 |
| MF (Amygdala) left | 53 | 56 | 0.01 | 0.00 | 0.01 | 1.00 | 0.00 | 0.99 |
| MF (Amygdala) right | 42 | 36 | 0.00 | 0.00 | 0.00 | 1.00 | 0.00 | 1.00 |
| SF (Amygdala) left | 39 | 41 | 0.06 | 0.06 | 0.00 | 1.00 | 0.06 | 0.94 |
| SF (Amygdala) right | 49 | 56 | 0.33 | 0.18 | 0.29 | 1.00 | 0.04 | 0.67 |
| Temporal to Parietal (GapMap) left | 55 | 66 | 0.19 | 0.10 | 0.19 | 1.00 | 0.00 | 0.81 |
| Temporal to Parietal (GapMap) right | 57 | 64 | 0.09 | 0.04 | 0.09 | 1.00 | 0.00 | 0.91 |
| VTM (Amygdala) left | 51 | 60 | 0.12 | 0.12 | 0.00 | 1.00 | 0.12 | 0.88 |
| VTM (Amygdala) right | 61 | 62 | 0.00 | 0.00 | 0.00 | 1.00 | 0.00 | 1.00 |
| Ventral Dentate Nucleus (Cerebellum) left | 62 | 65 | 0.16 | 0.08 | 0.16 | 1.00 | 0.00 | 0.84 |
| Ventral Dentate Nucleus (Cerebellum) right | 57 | 61 | 0.05 | 0.05 | 0.00 | 1.00 | 0.05 | 0.95 |
| Brainstem pons L (CSTL L + FPTL L)* | 46 | 57 | 0.45 | 0.22 | 0.45 | 0.00 | 0.00 | 0.55 |
| Brainstem pons R (CSTL R + FPTL R)* | 52 | 56 | 0.20 | 0.10 | 0.20 | 0.57 | 0.00 | 0.80 |

**Supplemental Figure 1**. Significant brain activations to pain stimulation were observed in all Julich atlas regions ($p < 0.05$, Bonferroni corrected) and in brainstem regions identified via a Human Connectome Project-based probability atlas.

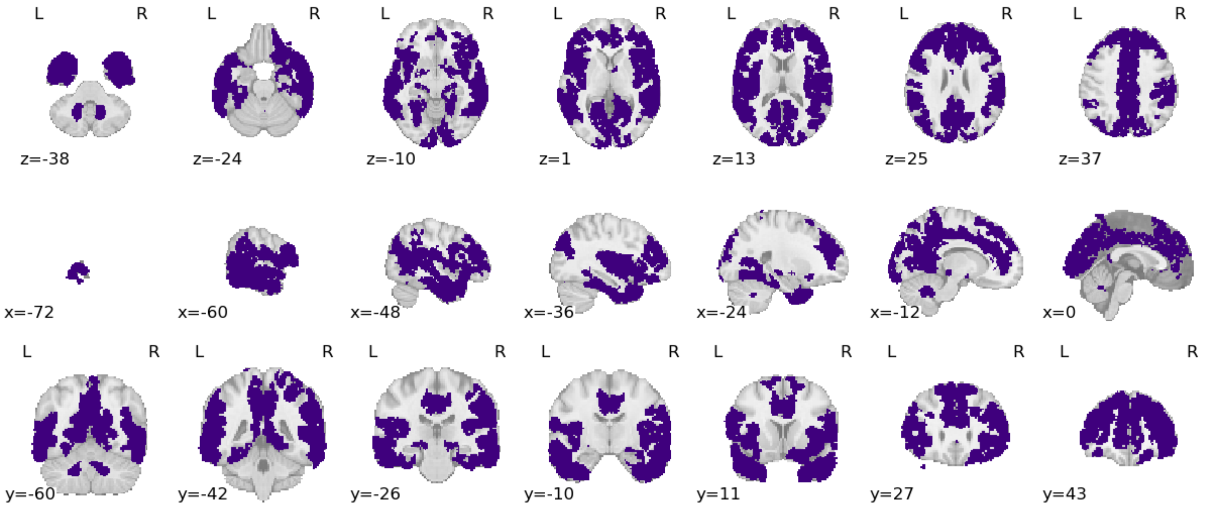
